# Plant microbiomes have a greater effect than plant size on flowering and growth in *Asclepias syriaca*

**DOI:** 10.64898/2026.10.20.759379

**Authors:** Joshua R. Puzey, Ivan Munkres, Christian D’Orgeix, Harmony J. Dalgleish

**Affiliations:** Biology Department, William & Mary, Williamsburg, VA 23185

## Abstract

Plants strike a delicate balance between growth and defense. The calculus of how to defend against herbivores, and maximize growth and reproduction, can differ between annuals and perennials. Annuals have a live fast, reproduce young strategy; while perennials may maximize fecundity over years. Milkweed, a herbaceous perennial, is a mix of these two strategies. On one hand, a single individual milkweed ramet is functionally a herbaceous annual and vulnerable to herbivory. On the other hand, below ground connections allow plants to share resources and persist for multiple years, perhaps lowering the cost of localized herbivory. Here, we study the impact herbivory on the leaf and soil microbiome on milkweed growth and flowering in a naturally occurring population. Specifically, using milkweed populations at three different sites in Virginia, following individual ramets throughout the growth and flowering season, we quantified the soil and leaf microbiome through metagenomic shotgun sequencing.

After controlling for plant size and plant genotype, we asked how the microbiome composition was related to flowering time and growth. We find that the leaf microbiome is a strong predictor of plant growth rate, but not flowering. Conversely, we find that the soil microbiome and plant size predict flowering probability. We found no evidence that herbivory altered the leaf or soil microbiome composition. Together these results provide evidence that plant microbiomes also influence growth and fecundity in natural populations of milkweed. These results also provide evidence that the leaf microbiome in milkweed is robust and stable in the face of significant herbivory. This study expands our knowledge of the contribution of plant microbiomes to plant demography in a clonal perennial plant.

## Introduction

Plants have evolved a myriad of defensive strategies to deter herbivores (Karban and Baldwin 2007; Agrawal 2007). Defense strategies are wide-ranging, including both chemical compounds and structural defense. While these defensive strategies are often effective at reducing or preventing herbivory, they often come with a significant energetic cost to the plant (Stamp 2003; Karasov et al. 2017). Plants must strike a balance between growth and fecundity while defending themselves against herbivores. Balancing the tradeoffs between growth and defense has long been recognized as a key component to optimizing plant fitness and fecundity (Herms and Mattson 1992).

While growth-defense tradeoffs in plants have been extensively studied, we have little understanding of how defensive compounds induced by herbivory may affect the plant-associated leaf and soil microbiomes. Microbiomes are important for plant growth and fecundity as they can alter nutrient uptake, pathogen defense, and plant stress tolerance (Bashir et al. 2022; Abadi et al. 2020; Mohanram and Kumar 2019; Xu et al. 2018). Plant chemical defenses against herbivory, can act as anti-bacterial and antifungal compounds, and change the microbiome of plants (Humphry and Whiteman 2020). Here, we address the question of whether herbivory changes leaf and soil microbiomes in ways that also affect plant growth and fecundity.

To explore the interaction of herbivory and microbial composition on plant growth, we use common milkweed (*Asclepias syriaca*) as a model system because the effects of herbivory on demography (growth and probability of flowering) are well established as are the growth defense trade-offs of cardenolides. For example, our previous research in milkweed shows that increasing herbivory severity decreases individual plant growth, survival, and fecundity, effects that we observe for both natural herbivory and experimental tissue removal (Dalgleish et al. 2024). Züst et al. (2015) found a negative correlation between cardenolide concentration and plant relative growth rate, supporting a tradeoff between growth and herbivory defense.

Increased cardenolide concentration also decreases fecundity and population growth (Struckman et al. 2019). Furthermore, several studies have shown that cardenolides have antibacterial properties against both Gram positive and Gram negative bacteria, fungistat activity, and anti-protozoan activity (Assob et al. 2011; Akhtar et al. 1992; Arora and Sood 2017; Jacobsohn and Jacobsohn 1985; Lefèvre et al. 2010), indicating that cardenolide concentration in leaves, which is both constitutive and induced by herbivory, may alter the composition of the plant’s leaf microbiome. If cardenolides leach into the rhizosphere, aboveground herbivory may also change the soil microbiome. Herbivory, then, may change plant growth, fecundity and the microbiome composition of the leaves and/or the soil.

Here, we use a field experiment to test how herbivory, the soil microbiome, the leaf microbiome, and plant genotype affect plant demography (growth and probability of flowering). To do this, we apply three levels of simulated leaf herbivory to milkweed plants across three populations of milkweed and measure plant growth and flowering during one season (Dalgleish et al. 2024). We determined the microbiome composition of the leaves and soil through metagenomic shotgun sequencing. Plant genotype was determined through low-pass sequencing. While we have shown in previous studies that herbivory decreases growth and reproduction (Struckman et al. 2019, Dalgleish et al. 2024), here we ask two additional questions:

1. Does herbivory alter the leaf and/or the soil microbiome?

We hypothesize that herbivory will affect leaf microbiome composition, with higher severity herbivory having the greatest change. Herbivory will affect the soil microbiome composition if herbivory alters the cardenolide concentration in the soil (Figure 1).

2. Does the leaf or soil microbiome influence plant growth and reproduction?

**Figure 1.**
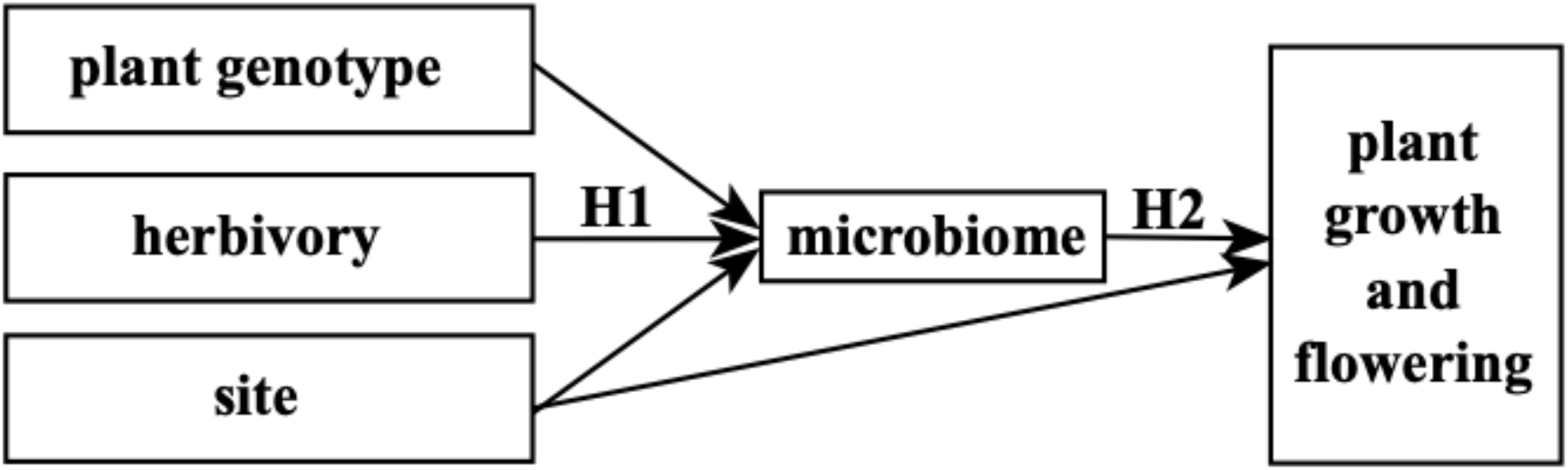
Conceptual diagram for our two main hypotheses. Hypothesis 1: Herbivory alters the plant microbiomes. Hypothesis 2: The microbiome will influence plant growth and reproduction independent of plant size. Plant genotype and site are considered as random effects in our analyses.

We hypothesize that both leaf and soil microbiomes will influence plant growth and reproduction independently of plant size (Figure 1).

## METHODS

For this study, we used data collected in 2021 from plants from two populations at Blandy Experimental Farm (39◦03′45.4′′N, 78◦03′46.9′W) and one population at Sky Meadows State Park (38◦59′11.1′′N 77◦57′41.6′′W). Complete site descriptions, plant selection protocol, herbivory treatments and field measurements are described in Dalgleish et al. (2024). Briefly, within each population we tagged 40 individuals and randomly assigned them to treatment groups that were subjected to artificial herbivory using a hole punch over six days. In the “low”, “medium” and “high” herbivory treatments respectively, approximately 5%, 25% and 75% of the leaf area was removed by the sixth day. We also had a control group that did not experience any hole punch damage. Because all plants experienced natural levels of herbivory over the course of the season, found that the low and no damage groups did not have significantly different amounts of tissue removed by the end of the season and, thus, we grouped the no damage plants with the “low” damage group for all analyses (suppl. Figure 1).

### Field measurements and sampling

The growth rate of each plant was calculated as the change in height divided by the number of days between measurements. Approximately 100 g of soil was collected with a trowel from the rhizosphere, 2-4 cm below the soil line and 2-10 cm away from the base of the plant. Soil samples were kept on ice during transport until frozen at-80°C. All soil samples were sieved through a double-mesh strainer to remove rocks and excess plant material before being processed for cardenolide concentrations or DNA extraction (see below). Leaf samples were taken near the apex after the completion of the treatments, placed on ice for transport, and stored at-80°C. In late June and early July, we recorded the number of inflorescences on each plant and we recorded seed pod production in September.

### Soil cardenolide quantification

Soil cardenolide concentrations were quantified for one site due to budget constraints. Sub-samples of rhizosphere soil were lyophilized in the dark for 24 hours or until dry. Then, 20 g of dry soil per sample was immersed in 100 mL of molecular-grade methanol, sonicated for 10 minutes in warm water, and centrifuged at maximum speed for 10 minutes. The supernatant was vacuum-filtered to remove soil solids. The samples were dried using a nitrogen dry-down tree, and 1 mL of molecular-grade acetonitrile was added. After sonication for 10 minutes, a 10 μg/mL digitoxin internal standard was added, and the samples were filtered through a 0.2 μm filter into a vial. Cardenolides were quantified using high-performance liquid chromatography (HPLC) with a photodiode array detector and a LiChrospher 100 RP-18 (5 μm) 250 mm x 4 mm HPLC column (Merck KGaA, Darmstadt, Germany) with a guard column, following the methods presented by Wiegrebe and Wichtl (1993). A 5 μL sample was injected, and the elution was performed at a flow rate of 1 mL per minute over a gradient of 84% to 5% acetonitrile in water for 30 minutes. Cardenolides were detected as symmetrical peaks between 205 and 235 nm, with λ_max_ between 214 and 224 nm. The average gross cardenolide concentration was calculated for each sample using the combined spectral areas. Multiple regression analysis was used to assess the effects of average herbivory score, plant size (represented by height), and location within the site on soil cardenolide concentration.

### DNA extraction and sequencing

We extracted DNA from soil samples using a FastDNATM Spin Kit for Soil (MP Biomedicals LLC, Irvine, CA) using 500 mg of soil per sample. We cleaned the extracted DNA by using a DNA Clean & Concentrator® Kit (Zymo Research, Irvine, CA); we then treated with RNAse. We extracted DNA from leaf samples using a DNeasy® Plant Mini Kit (QIAGEN, Germantown, MD). The extracted DNA was cleaned and concentrated in the same manner as the soil samples and were also treated with RNAse. Multiplexed DNA libraries were prepared using plexWellTM 96-well DNA Library Prep Kits (seqWellTM, Beverly, MA) for both soil and leaf DNA samples. The libraries were sequenced on an Illumina NovaSeq 6000 SP sequencer by the Michigan State University RTSF Genomics Core.

### Plant genomic analyses

We aligned the DNA to the milkweed genome (Boyle et al. 2023) using bowtie2 (Langmead et al. 2009). Following alignment, GATK HaplotypeCaller was used to call genotypes for each individual producing a VCF file (McKenna et al. 2010). The VCF file for each individual was filtered to remove sites not supported by a minimum depth of 10 using VCFtools and then merged into a composite VCF file. Next, using VCFtools, the VCF file was filtered using these parameters (--max-missing 0.5 --min-meanDP 2 --minQ 30) and then filtered for linkage using Plink (indep-pairwise 50 10 0.1) (Purcell et al. 2007). A PCA of the resultant filtered genotypes was conducted using Plink.

### Microbiome data analysis

We used Metalign, a tool developed for analysis of metagenomic data, to analyze the microbiome data (LaPierre et al. 2020). Prior to analysis with Metalign, we aligned the sequencing reads to the milkweed reference nuclear genome, chloroplast genome, and mitochondrial genome (Boyle et al. 2023). Reads that mapped to these plant genomes were filtered from the raw sequencing data prior to analysis with Metalign.

Metalign allows for characterization of results at different taxonomic levels. Using the Metalign results, we analyzed the soil and leaf microbiome data at the ‘family’ level.

Microbiome taxa were removed when more than 80% of all samples were missing presence data. For the remaining data, NA values were converted to zero. Both datasets were scaled using z-transformation and a PCA was performed. The family composition of the leaf microbiome was 168 Bacteria, 31 Eukaroyta, four Archaea, and four viruses. The family composition of the soil microbiome was 341 Bacteria, one Eukaroyta, 14 Archaea, and four viruses.

## RESULTS

### Sites explains a small amount of the variation among plant genotype and microbiomes

We found some evidence of milkweed genetic differentiation by site. A subset of the population at Blandy Meadow separated along the first plant genotype PC axis, while the other half of Blandy Meadow had broad genetic overlap with both Sky Meadows and Blandy Village (suppl. Figure 2). We also found evidence that microbiomes were different among sites, separating along the first PC axis for both the soil and leaf microbiomes (suppl. Fig. 2). For both microbiomes, Blandy Village differed from both Sky Meadows and Blandy Meadow (suppl. Fig. 2). Plant genotype was not related to either the soil microbiome (*P* = 0.09) or the leaf microbiome (*P* = 0.62; suppl. Figure 3). In addition, the first PC axes of the soil and leaf microbiomes were not related to each other (*P* = 0.89; suppl. Fig 3). While we found a significant relationship between soil microbiome PC2 and leaf microbiome PC1, this relationship was entirely driven by site effects (suppl. Figure 4; *P* = 0.22 when site is included as a random effect). In all subsequent analyses, we included site as a random effect in the statistical models when it explained significant variation.

### Soil cardenolide concentrations are related to plant genotype, but not plant size or herbivory

Cardenolides were detected in all of the 26 soils samples we analyzed and varied considerably among plants with an average concentration of 0.372 ± 0.281 μg/g (mean ± SD). Soil cardenolide concentration was not different among herbivory categories (suppl. Fig 5). We also found no effect of stem diameter or plant height on soil cardenolide concentrations (suppl. Fig 5). Plant genetic identity, however, was related to soil cardenolides (*ρ* = 0.58; *P* = 0.03, suppl. Fig. 6).

### The leaf microbiome is the best predictor of plant growth rate

First, we modeled plant growth rate as a function of plant genotype (PC1), soil microbiome (PC1), leaf microbiome (PC1), and herbivory category in order to determine which of these predictors contributed to plant growth (n = 49). Site did not explain significant variation and was dropped as a random effect. Because the leaf microbiome PC1 is strongly correlated with plant height (*ρ* =-0.72, *P* < 0.001) and diameter (*ρ* =-0.65, *P* < 0.001; suppl. Fig. 5), we did not include a measure of plant size in this initial growth model. The p-value for leaf microbiome (*P* < 0.001) and herbivory category (*P* = 0.067) indicated that these variables may be important for predicting plant growth. None of the other variables had low *P*-values (all were greater than 0.12).

We then analyzed the leaf and soil microbiome datasets separately, which increased our sample size from 49 to 63 plants (i.e., 63 plants had leaf data but only 49 had leaf and soil data). We modeled growth as a linear function of either the soil microbiome (PC1) or the leaf microbiome (PC1) and herbivory category (Figure 2). Herbivory was not significant in either model, the soil microbiome was not significant, but leaf microbiome remained significant (*P* < 0.001).

**Figure 2.**
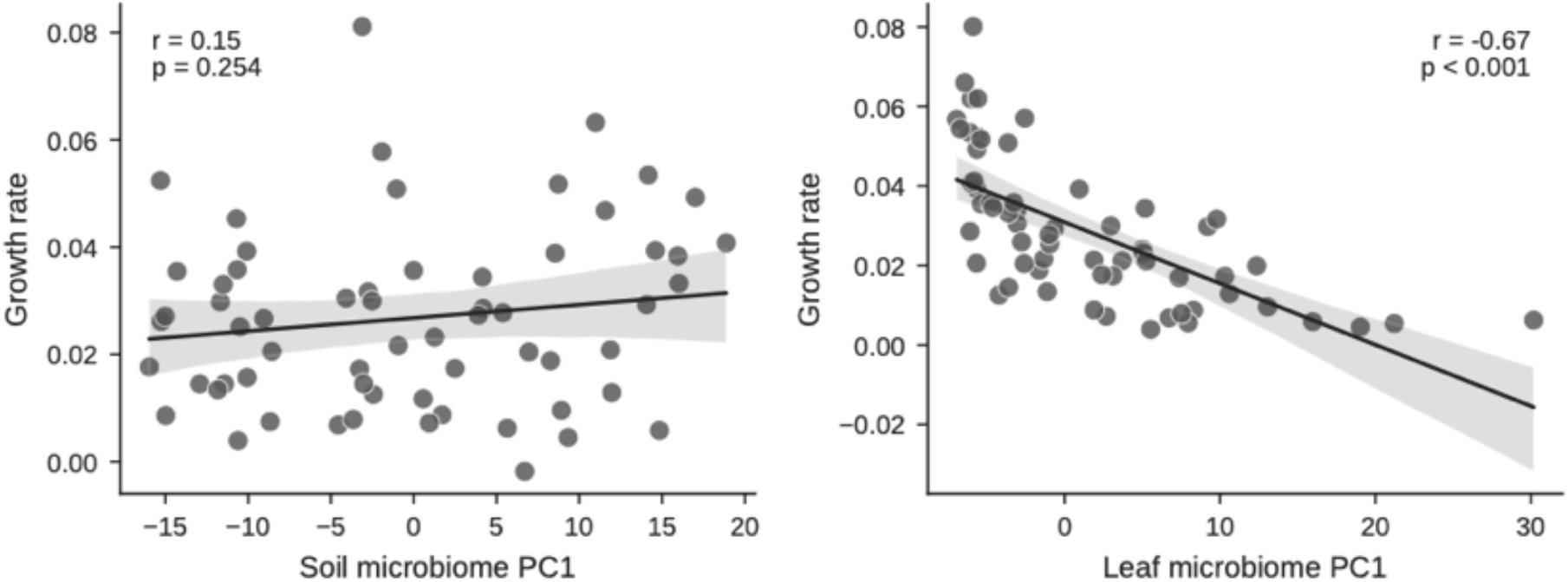
The effect of the plant microbiome on growth rate. Soil microbiome composition (PC1) does not affect plant growth rate while leaf microbiome composition (PC1) is negatively related to plant growth (n = 62)

To examine the effects of plant size and the leaf microbiome on plant growth, we first fit a full model with both predictors, which explained a substantial portion of the variation in plant growth rate (*r^2^* = 48.9%). Variance partitioning revealed that the majority of this explained variation was attributable to the leaf microbiome (LMG = 30.0%), while plant size explained less (LMG = 18.9%). Finally, because the leaf microbiome and plant size are correlated (*ρ* = - 0.65), to isolate the effects of plant size and the leaf microbiome on plant growth, we extracted the residuals from a linear model where plant size predicted leaf microbiome PC1 (leaf residuals) and a separate linear model where the leaf microbiome PC1 predicted plant size (size residuals). With the effects of plant size removed, the leaf microbiome remained a significant predictor of plant growth (*F* = 10.34_1,60_, *P* = 0.002, *r^2^* = 0.14). Plant size, however, did not remain a significant predictor of plant growth when we removed the effects of the leaf microbiome (*F* = 2.27_1,60_, *P* = 0.14, *r^2^* = 0.04).

### The soil microbiome and plant size predict the probability of flowering

We followed a similar approach to modeling the probability of flowering with plant genotype (PC1), soil microbiome (PC1), leaf microbiome (PC1), and herbivory category as predictors. Site did not explain significant variation and was dropped as a random effect. None of these predictors were significant. We then analyzed the soil and leaf microbiome datasets separately, which increased our sample size from 49 to 63 plants (i.e., 63 plants had leaf data but only 49 had leaf and soil data). We modeled the probability of flowering as a generalized linear model of either the soil microbiome (PC1) or the leaf microbiome (PC1) and herbivory category. Herbivory was not significant in either model.

The leaf microbiome was a significant predictor of the probability of flowering as was plant size (stem diameter). Because the leaf microbiome and plant size are correlated (rho = - 0.65), to isolate the effects of plant size and the leaf microbiome on plant growth, we extracted the residuals from a linear model where plant size predicted leaf microbiome PC1 (leaf residuals) and a separate linear model where the leaf microbiome PC1 predicted plant size (size residuals). With the effects of plant size removed, the leaf microbiome was no longer a significant predictor of the probability of flowering (deviance = 0.14; *P* = 0.70) (suppl. Fig. 7). Plant size, however, did remain a significant predictor of the probability of flowering when we removed the effects of the leaf microbiome (deviance = 59.1; *P* < 0.001).

The soil microbiome (PC1; deviance 17.7, *P* < 0.001) and plant size (stem diameter; deviance = 5.0, *P* = 0.03) were both significant predictors of the probability of flowering (Figure 3). SoilPC1 is significant with soil odds ratio per SD = 4.64, *P =* 0.015, and stem diameter OR per SD = 3.00, *P* = 0.05). A partial distance based redundancy analysis predicting euclidean distance of the leaf microbiome composition using herbivory rate and conditional variables of plant genotype and stem diameter determined that 1.07% of leaf microbiome variation is explained by herbivory.

**Figure 3.**
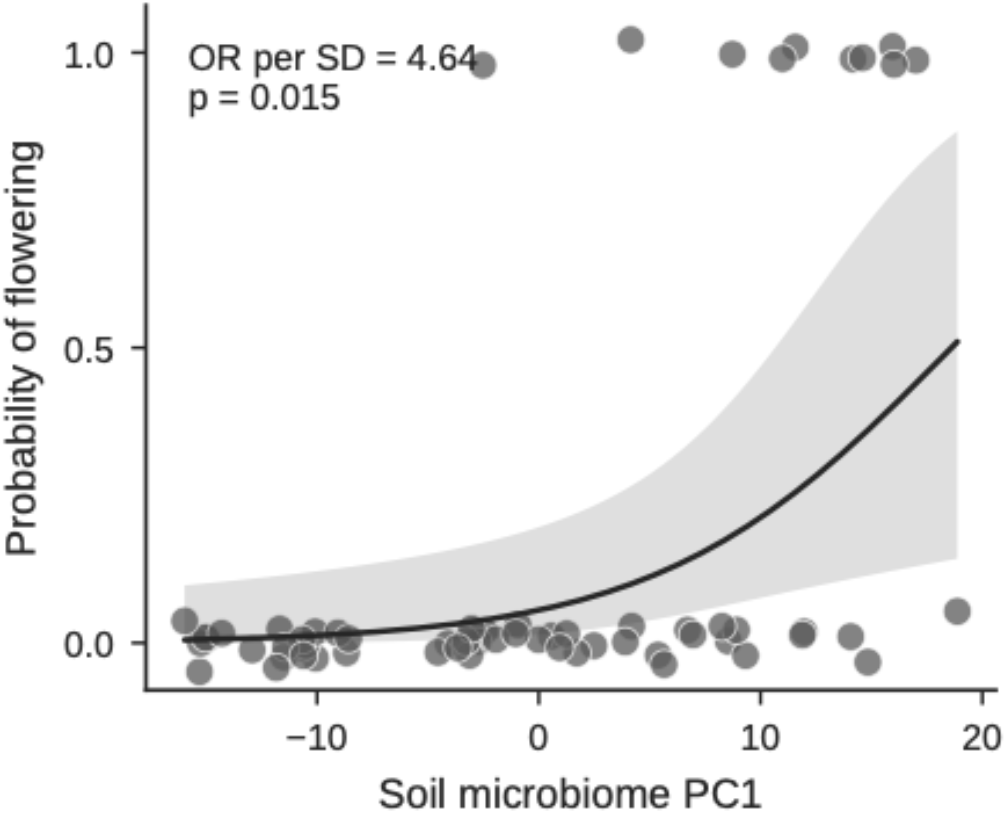
Soil microbiome PC1 alters plant flowering after accounting for stem diameter. Soil PC1 and stem diameter both show significant effects on probability of flowering (Soil PC1 odds ratio per SD = 4.64, *P* = 0.015, n = 62; Stem diameter odds ratio per SD = 3.00, *P* = 0.050, n = 62).

## DISCUSSION

Traditionally, plant size has been considered the main driver of plant demography. Here we provide evidence that plant microbiomes also influence growth and fecundity. However, they affect different aspects of plant demography: the leaf microbiome influenced plant growth while the soil microbiome, combined with plant size, influenced the probability of flowering. We found no evidence that herbivory severity directly altered either growth or the probability of flowering, nor did it change microbiome composition. Rather, herbivory likely affects plant demography through its effects on plant size, which, combined with the soil microbiome composition influences the probability of flowering.

Relatively few studies have examined the impact of the leaf microbiome on flowering. What is known is about the relationship of the leaf microbiome to flowering comes primarily from studies in annual plants grown in laboratory settings. For instance, experimental manipulation of leaf microbial communities in Arabidopsis has been shown to alter flowering and reproduction (Saleem et al. 2017; Zahn and Amend 2019). Similarly, using Solanum as a model system, Mehlferber et al. found that a synthetic leaf microbiome altered fruit production for plants grown in greenhouse settings (Mehlferber et al. 2023). While important research, these studies had similarities in that they all manipulated the leaf microbiome, grew plants in artificial settings, and conducted studies using short lived annuals as model systems. To our knowledge, no studies have looked at the relationship of the leaf microbiome on flowering in the natural settings or in longer lived perennials. Our study begins to fill that gap. In milkweed grown in natural settings we found no evidence that the leaf microbiome composition was related to flowering probability. Plant size is a strong predictor of flowering probability in milkweed (Dalgleish et al. 2024). This fact, combined with the conclusion the milkweed leaf microbiome impacts plant growth, we expected to find an indirect effect of the leaf microbiome on plant flowering. However, we were not able to detect this effect in our dataset. Why is this the case and how do we make sense of this result? In milkweed, the lack of a clear relationship between leaf microbiome composition and flowering probability likely indicates a more complex relationship between the phyllosphere microbiome, plant size and flowering probability. Whether this result stands up to experimental manipulation and is the case in other longer-lived herbaceous perennials is an exciting and open question.

We also found no evidence that herbivory altered the milkweed leaf microbiome composition. This result is in contrast to other studies which found that herbivory alters the leaf microbiome. For example, in Camellia, it was found that herbivory reduced microbial diversity (Wang et al. 2026). In Cardamine and Arabidopsis, herbivory was shown to alter the leaf microbiome composition (Humphrey and Whiteman 2020; Müller et al. 2026). There are several reasons for these different results – both methodological and biological. First, we recognize that the lack of herbivory effect could be methodological. Our study differs from these other studies in that we use metagenomic shotgun sequencing instead of 16S sequencing to quantify the microbiome. While each method has issues, calculating microbial composition using 16S is difficult with many well recognized sources of bias (Pollock et al. 2018; Brooks et al. 2015).

While metagenomic shotgun sequencing is not perfect, it can provide a better estimate of microbial composition (Ranjan et al. 2016). It is possible that the use of metagenomic shotgun sequencing in this study provided a less biased estimate of microbiome composition leading to different results. Another possibility is that there was an herbivory effect but we were unable to detect it, perhaps due to the fact that our response was overall composition (leaf microbiome PCA) and not individual taxa. Or finally, the lack of herbivory effect could be pointing to a real biological signal and indicate that milkweed has a more stable leaf microbial community in the face of significant herbivory. Mechanistically, in plants, we expect the leaf microbial community to be altered due to change in nutrient and carbon resources available to the microbes. In milkweed, perhaps the resource shifts due to herbivory are mitigated by the fact that plants are clonal with underground connections allowing for individual ramets to have more stable nutrient and carbon resources. Stable resources may lead to more stable microbiomes.

Although both plant size and the soil microbiome composition were significant predictors of the probability of flowering, the soil microbiome had a larger effect than plant size. The important role of the soil microbiome in plant life history, particularly flowering, is only recently coming to light (O’Brien et al. 2021; Aizaz et al. 2025). Many studies of annuals and short-lived perennials demonstrate that different soil microbiomes can shift flowering phenology, either earlier or later (Wagner et al. 2014; Panke-Buisse et al. 2015; Chaney and Baucom 2020), and can increase the number of flowers (Ramula et al. 2025; Chaney and Baucom 2020). In one of the few studies of a perennial, Ramula et al. (2025) found the presence of the natural microbial community increased plant growth and increased the population growth rate (lambda) estimated from an integral projection model (IPM) based upon plant size. However, their analysis did not disentangle the effects of the microbiome from plant size as we do here. Using an IPM with the microbial community in addition to plant size is an excellent next step to understand how plant-microbe interactions affect plant populations. While our observational study cannot tease apart the mechanism of how microbiomes affect flowering in milkweed, we hypothesize that it relates to different nutrient availability with different microbial communities. Microbes that change nutrient availability, are pathogenic or change phytohormones have all been associated with increased or decreased flowering or shifted phenologies (O’Brien et al. 2021).

Plant secondary metabolites can alter the plant soil microbiome, particularly the microbes in the rhizosphere (Schütz et al. 2021). When plants respond to herbivory with induced defenses, these can alter the soil microbiome (Hu et al. 2018). Although we found cardenolides in the soil, the concentration was not related to the herbivory treatment. This may explain why we found no effect of herbivory on the soil microbiome. However, plant genetic identity was related to soil cardenolide concentration, although genetic identity did not correlate with either soil or leaf microbiomes. Cardenolides, then, may not play a role in structuring milkweed microbiomes. One caveat is that we quantified the soil microbiome using PCA. While this allows us to look at overall community changes, there may be differences in the identity of microbes that occur, or in the richness of rare taxa, that this method would mask.

Our study provides support for the idea that microbes are a silent third partner in species interactions. Importantly, this study expands our knowledge of plant microbiome interactions in two important ways: both in the direction of a field based demographic study and by expanding our understanding to a clonal perennial plant. In the future, a manipulative experiment to test the results here and ask whether the patterns found here extend to other milkweed species would shed light on the broader generalizability of these results. Also a study focused on disentangling the effect of clonal below ground connection on the microbial community could be a particularly ripe area of research.

## Supplemental Figures

**Supplemental Figure 1:**
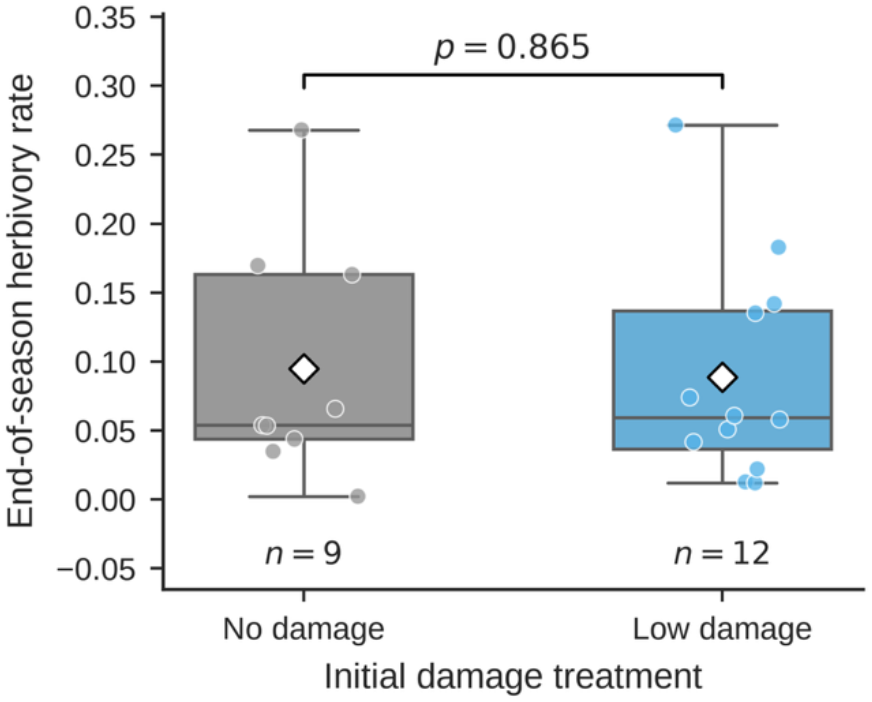
“No damage” plants were not different from “low damage” plants in end of season herbivory level due to natural occurring herbivory.

**Supplemental Figure 2:**
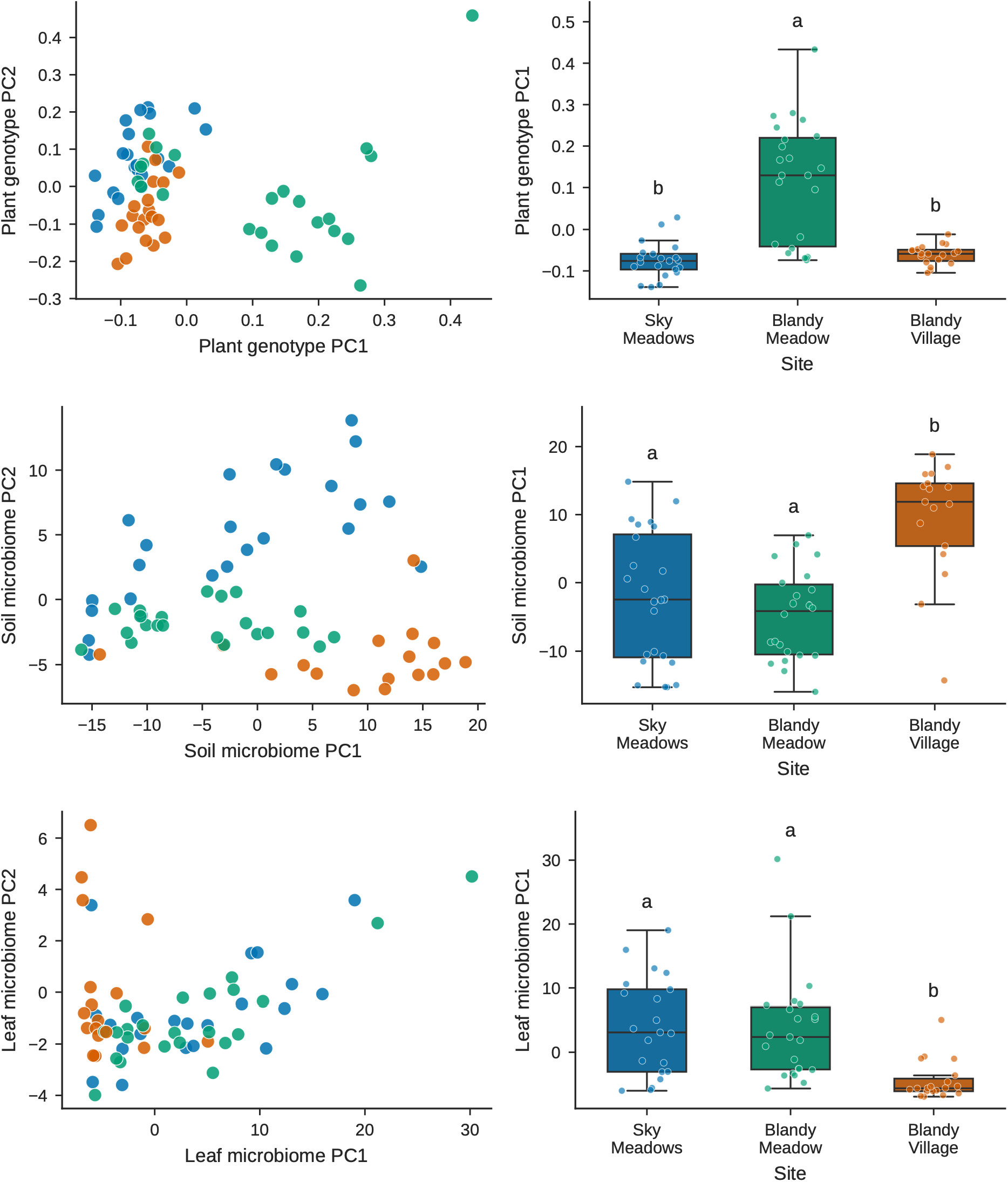
Relationship of site to plant genotype, soil microbiome, and leaf microbiome

**Supplemental Figure 3:**
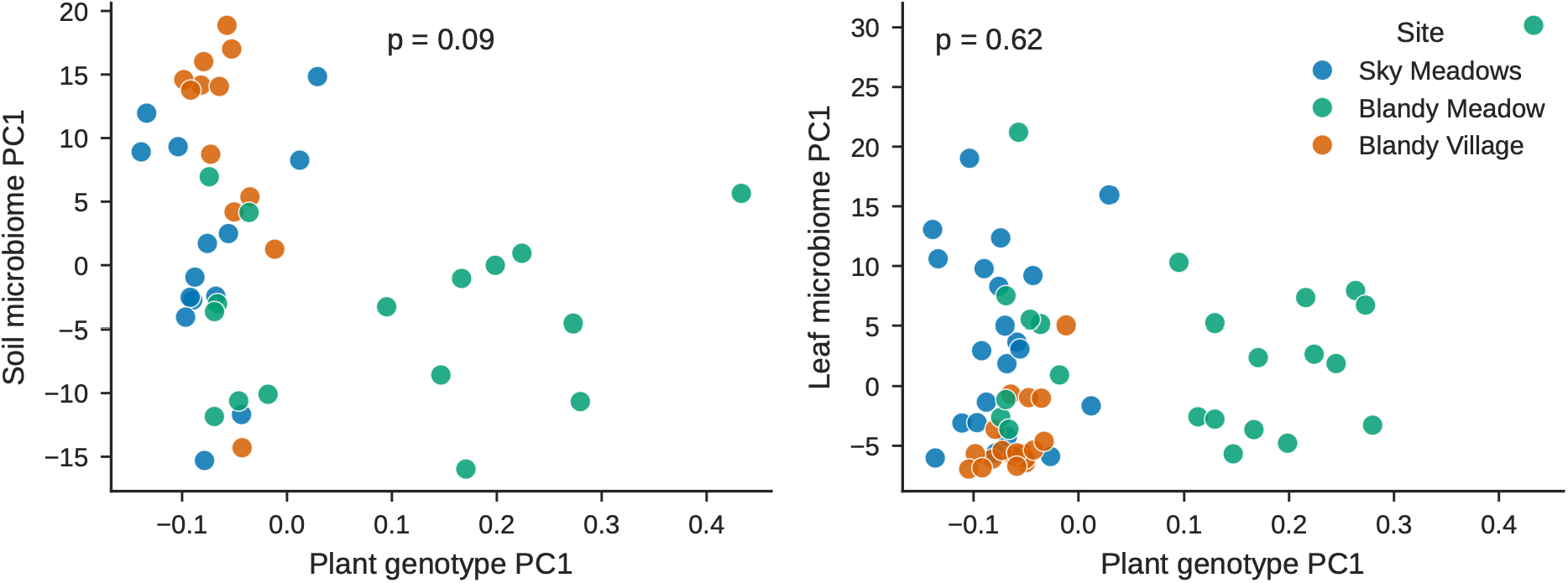
Relationship of plant genotype to soil and leaf microbiome

**Supplemental Figure 4:**
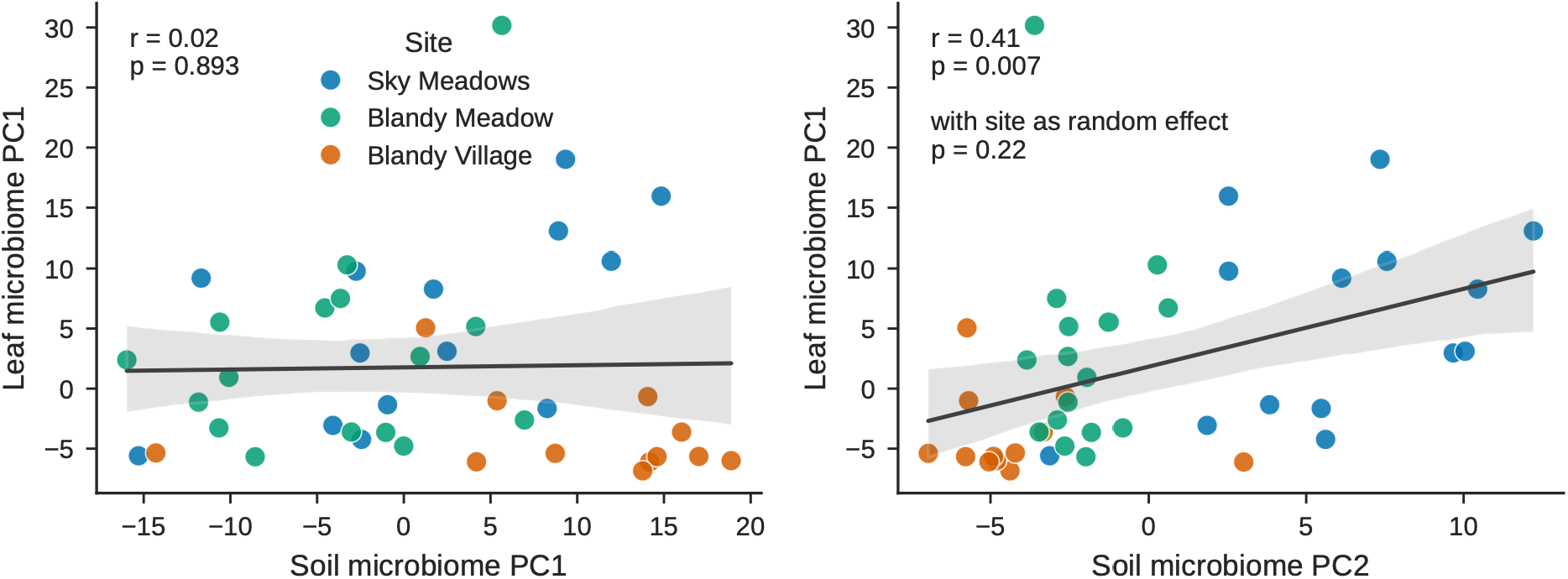
**Relationship of the soil and leaf microbiome composition**

**Supplemental Figure 5:**
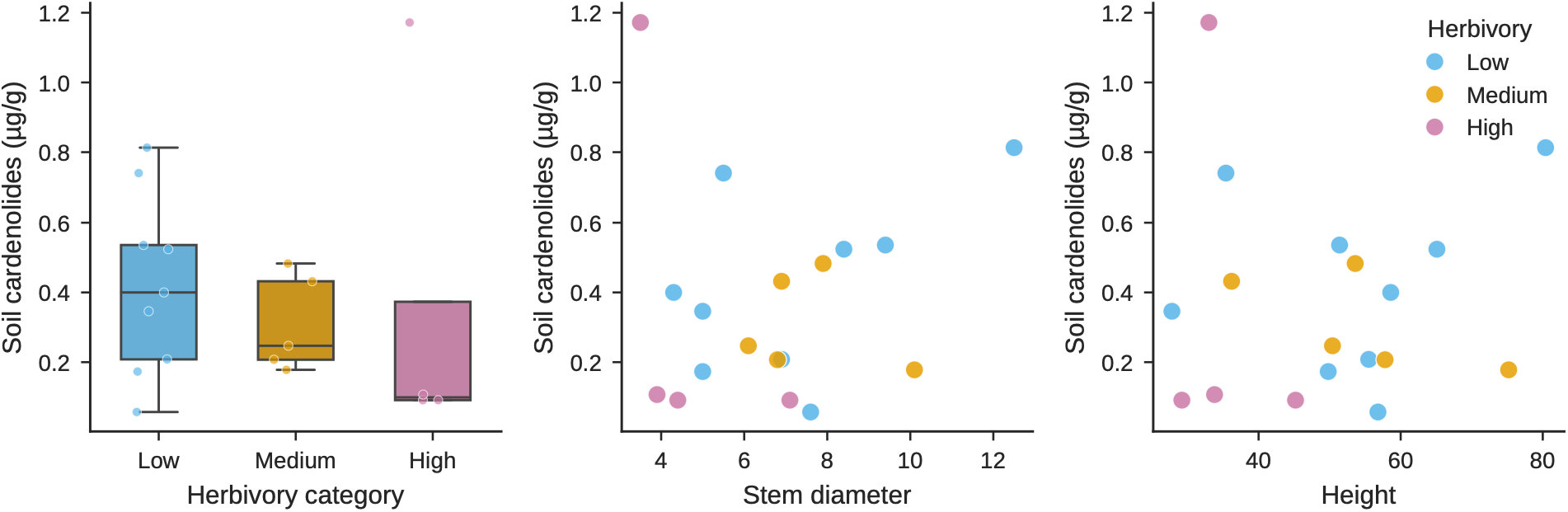
Effect of herbivory and plant size on soil cardenolide concentration. Analysis was conducted with data as shown and no effect of herbivory or plant size was detected (p>0.05). Analysis was also conducted with outlier ‘high-herbivory’ cardenolide removed and results were same.

**Supplemental Figure 6:**
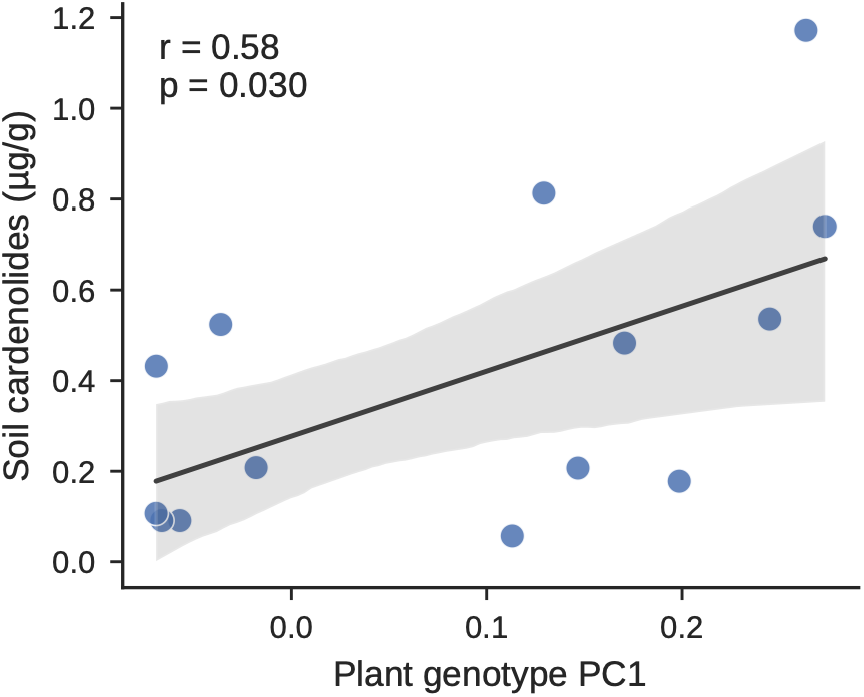
Plant genotype PC1 is positively related to soil cardenolide concentration.

**Supplemental Figure 7:**
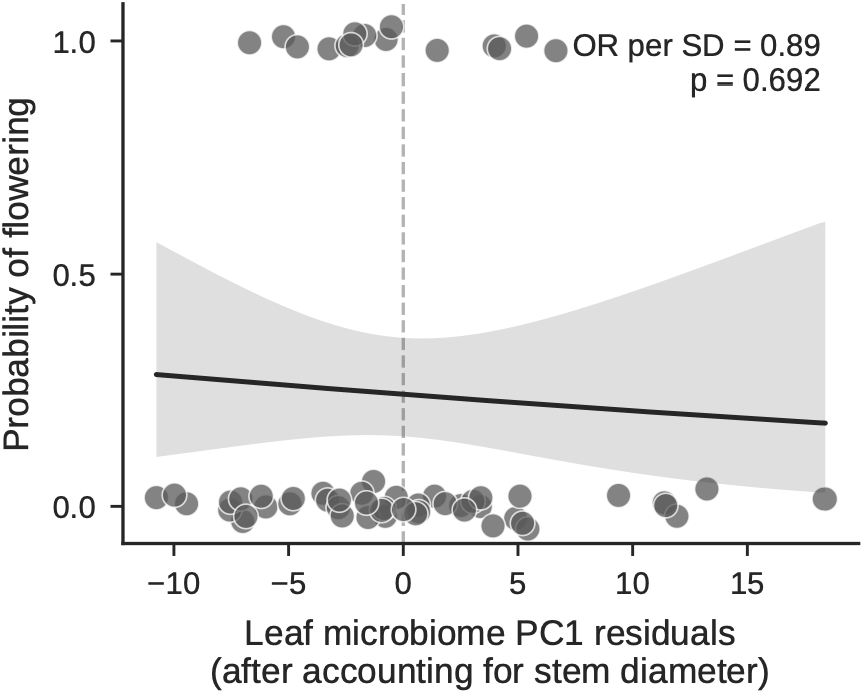
LeafPC1 residuals (after accounting for stem diameter) shows no effect on the probability of flowering

